# Dynamics of the most common pathogenic mtDNA variant m.3243A>G demonstrate frequency-dependency in blood and positive selection in the germline

**DOI:** 10.1101/2021.02.26.433045

**Authors:** Melissa Franco, Sarah J. Pickett, Zoe Fleischmann, Mark Khrapko, Auden Cote-L’Heureux, Dylan Aidlen, David Stein, Natasha Markuzon, Konstantin Popadin, Maxim Braverman, Dori C. Woods, Jonathan L. Tilly, Doug M. Turnbull, Konstantin Khrapko

## Abstract

The A-to-G point mutation at position 3243 in the human mitochondrial genome (m.3243A>G) is the most common pathogenic mtDNA variant responsible for disease in humans. It is widely accepted that m.3243A>G levels decrease in blood with age, and an age correction representing ∼2% annual decline is often applied to account for this change in mutation level. Here we report that recent data indicate the dynamics of m.3243A>G are more complex and depend on the mutation level in blood in a bi-phasic way. Consequently, the traditional 2% correction, which is adequate ‘on average’, creates opposite predictive biases at high and low mutation levels. Unbiased age correction is needed to circumvent these drawbacks of the standard model. We propose to eliminate both biases by using an approach where age correction depends on mutation level in a biphasic way to account for the dynamics of m.3243A>G in blood. The utility of this approach was further tested in estimating germline selection of m.3243A>G. The biphasic approach permitted us to uncover patterns consistent with the possibility of positive selection for m.3243A>G. Germline selection of m.3243A>G shows an ‘arching’ profile by which selection is positive at intermediate mutant fractions and declines at high and low mutant fractions. We conclude that use of this biphasic approach will greatly improve the accuracy of modelling changes in mtDNA mutation frequencies in the germline and in somatic cells during aging.

## Introduction

Pathogenic variants in the mitochondrial genome are responsible for a wide range of diseases that affect mitochondrial function (Gorman et al., 2016). The multi-copy nature of mitochondrial DNA (mtDNA) means that it is possible for more than one species of mtDNA to co-exist within the same cell, termed heteroplasmy. By far the most common heteroplasmic mtDNA pathogenic variant is an A to G transition at position 3243 (m.3243A>G) within MT-TL1, which encodes mitochondrial tRNA^Leu(UUR)^ (Goto et al., 1990).Estimates of m.3243A>G carrier frequency range from 140 to 250 people per 100,000 (Elliott et al., 2008; Manwaring et al., 2007), although the point prevalence for adult disease is much lower than this, at 3.5 per 100,000 (Gorman et al., 2015), suggesting that many carriers are either asymptomatic or have mild, undiagnosed symptoms.

Originally identified within a cohort of patients presenting with a severe syndrome characterized by mitochondrial encephalopathy, lactic acidosis and stroke-like episodes (MELAS), m.3243A>G is associated with extremely varied clinical presentations. Patients can experience a variety of phenotypes including ataxia, diabetes, deafness, ptosis, chronic progressive ophthalmoplegia, cardiomyopathy, cognitive dysfunction and severe psychiatric manifestations (de Laat et al., 2012; Fayssoil et al., 2017; Koga et al., 2000; Mancuso et al., 2014; Nesbitt et al., 2013; Pickett et al., 2018). Disease burden can be partly explained by an individual’s m.3243A>G mutation level, but this relationship is not simple; other factors are likely to also play a role (Boggan et al., 2019; Grady et al., 2018; Pickett et al., 2018). These complexities make offering prognostic advice to patients very difficult and, coupled with the mutation’s high frequency, mean that m.3243A>G-related disease is one of the biggest challenges in the mitochondrial disease clinic.

Although levels of m.3243A>G are relatively stable in post-mitotic tissues such as muscle, there is strong evidence of negative selection against the G allele in mitotic tissues, even in relatively asymptomatic patients; this loss of m.3243A>G in mitotic tissues has been studied in detail in blood (de Laat et al., 2012; Grady et al., 2018; Langdahl et al., 2018; Mehrazin et al., 2009; Pyle et al., 2007; Rajasimha et al., 2008; Sue et al., 1998). Initial simulation studies suggested that this decline could be exponential (Rajasimha et al., 2008), although more recent studies using larger amounts of data point to a more complex process (Grady et al., 2018; Veitia, 2018). To study the dynamics of this mutation, we took advantage of the large quantity of longitudinal data that are available for m.3243A>G levels in blood and have developed a new, empirical model that better describes the dichotomous pattern that we observe.

The m.3243A>G variant is maternally inherited and, similar to other heteroplasmic mtDNA mutations, undergoes a genetic bottleneck in development leading to offspring often with very different levels of m.3243A>G than their mothers (Chinnery et al., 2000; Pickett et al., 2019). Different mtDNA mutations segregate at different rates, demonstrating that the dynamics of this bottleneck are dependent on the mtDNA variant being transmitted (Wilson et al., 2016). We postulated that studying this bottleneck may help us understand why the m.3243A>G variant is so common in the population; using our new model of the dynamics of blood heteroplasmy/mutation level, we explored whether there is a transmission bias for m.3243A>G between mother and child, i.e. whether m.3243A>G is under germline selection.

## Results

### Longitudinal changes of m.3243A>G levels in blood with age reveal a dichotomous pattern (Figure 1)

A large amount of data on the changes of the level of m.3243A>G mutation in blood with the age of the individual have been compiled in a recent report (Grady et al., 2018). We have reproduced these data in Figure 1A. Traditionally, the dynamics of m.3243A>G mutation in the blood are considered to follow an exponential decay with age of approximately 2% per year. The data in Figure 1A, however, appear visually dichotomous, i.e., at higher m.3243A>G levels we see more cases where the mutation level decreases (red), whereas at lower levels there is little evidence of decline and there are more cases with an increase or that remain constant (blue). The dichotomy can be further demonstrated by binary logistic regression analysis (**Figure 1B**). Figure 1B shows that individuals with higher m.3243A>G levels tend to, in accordance with the conventional model, decrease their levels with age, while contradictory to the convention, individuals with low levels tend to increase or stabilize their mutation level. This trend is significantly non-random (the slope of regression curve is significantly negative p<0.0001).

**Fig. 1.**
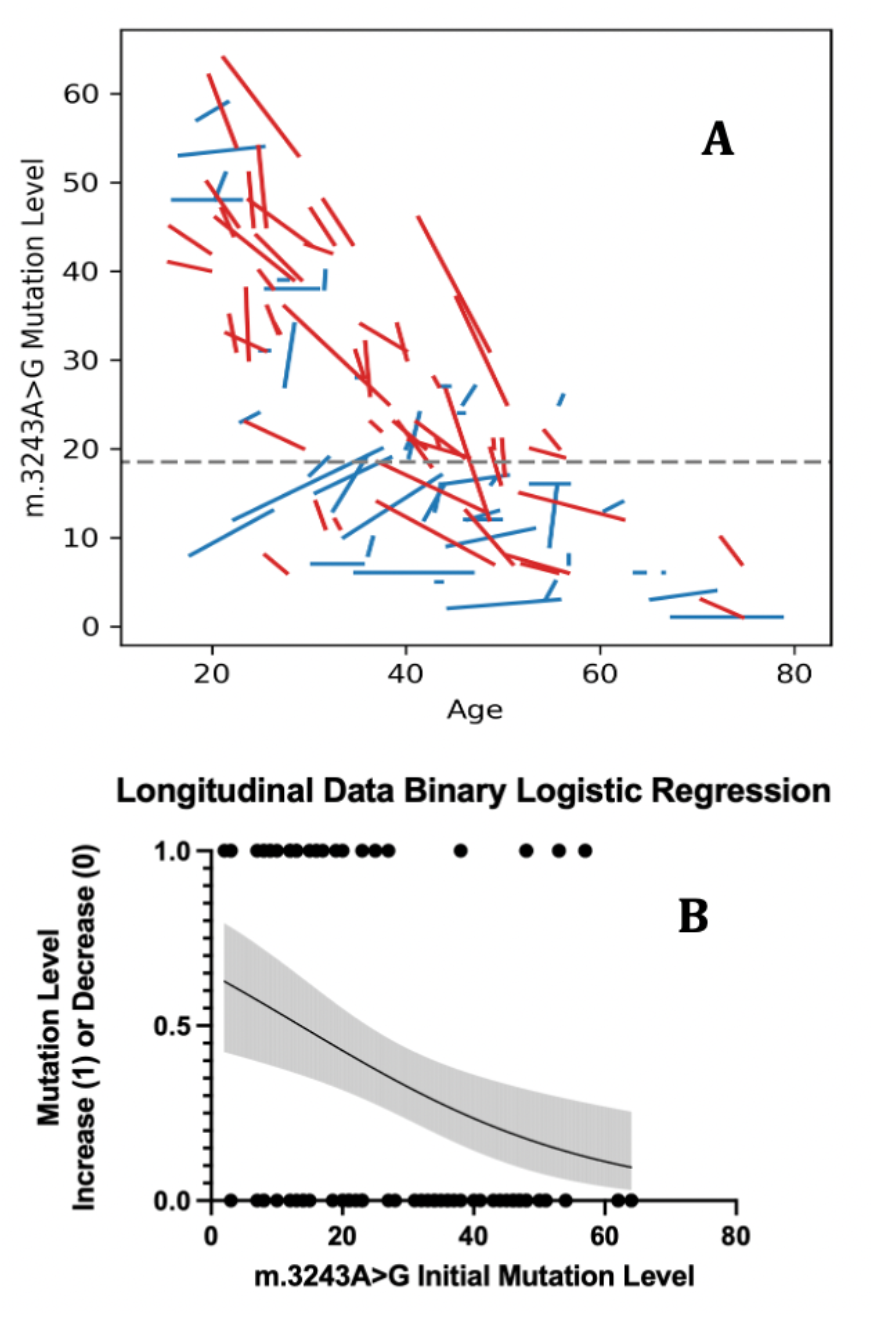
Analysis of longitudinal of m.3243A>G levels in blood. (**A**) Each segment represents the change in m.3243A>G level for a single individual for the follow-up period. Red and blue lines represent cases of age-related increase or decrease/stable m.3243A>G levels, respectively. **(B**) Logistic regression of the data presented in A. Increasing segments are represented as ‘1s’, and decreasing as ‘0s’. Logistic regression curve has been constructed and significance of the negativity of the curve estimated (P<0.0001). (See Materials and Methods for details).

### Mother-child m.3243A>G levels also show a dichotomous pattern

The dichotomous pattern of m.3243A>G dynamics in blood is further supported by the analysis of a different though similarly constructed dataset, i.e. the mother-child dataset (Pickett et al., 2019), that represents inheritance of m.3243A>G between mothers and their children. We note that this dataset shows a similar dichotomous pattern (Figure 2A) as the longitudinal dataset, which reflects the dynamics in blood with age discussed above (Figure 1A). Indeed, in the high child mutation level range (approximately above 5%) the majority of mother/child relationships are strongly descending in mutation frequency. In contrast, in the low child mutation level region (below 5%), the ascending child>mother pattern is prevailing. In accordance with this visual appearance, logistic regression analysis confirms, as with longitudinal data in Figure 1B, a statistically significant increase of child-mother pairs with increasing mutation levels (blue segments) at lower child mutant fractions. We conclude that in both the longitudinal and the mother/child datasets, increase of m.3243A>G mutation level prevails at the low mutation levels and decrease prevails at the high mutation levels. Thus, the rate and the direction of change of m.3243A>G level in blood appear to depend on the mutation level.

**Fig. 2:**
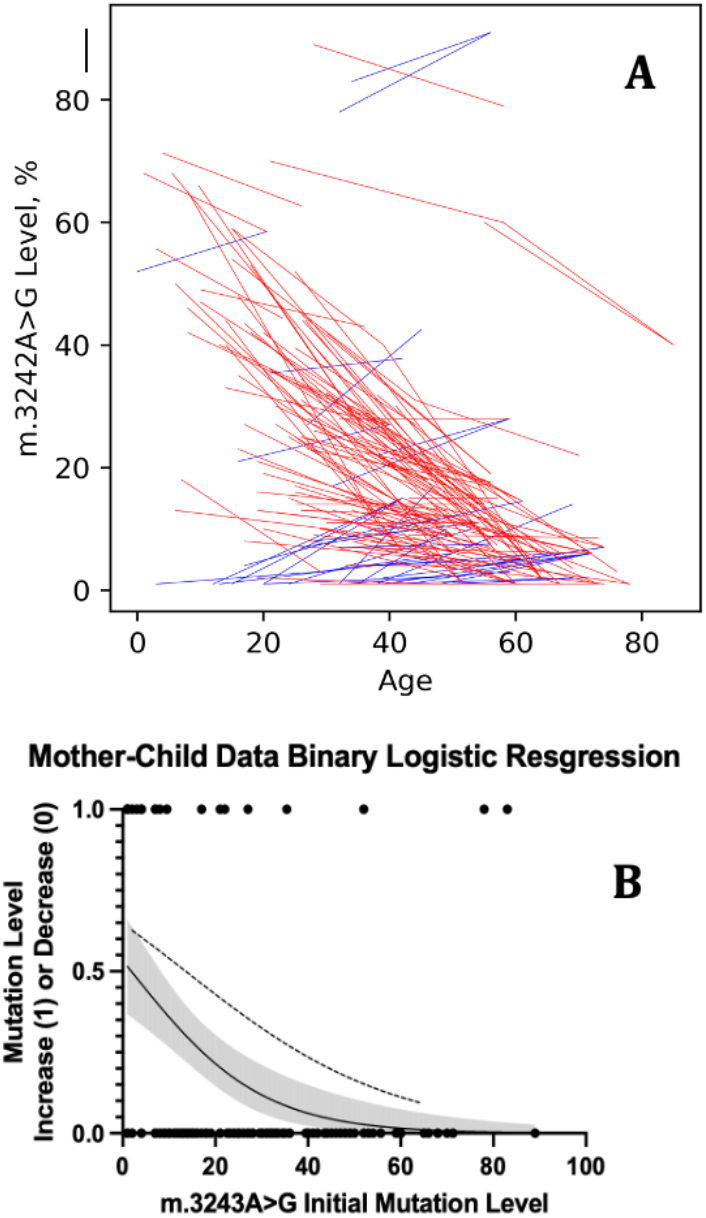
Analysis of mother-child m.3243A>G levels in blood. (**A**) Every segment connects the m.3243A>G level of a child (left end of the segment) to the m.3243A>G level of their mothers (right end). Descending segments are coded red, ascending - blue. (**B**) Solid curve: Logistic regression curve of the mother-child data presented in A, constructed as in Fig 1B. Broken curve: longitudinal data logistic curve from Fig 1B, overlaid for comparison. The grey shaded curve represents the 95% confidence interval for the binary regression at each point of the curve.

### The standard 2% annual decline model is biased at high and low mutational levels

The observed dichotomy of m.3243A>G (Figures 1 and 2) implies that, because the conventional 2% annual decline model inherently applies to all mutation frequency levels in the same way and thus cannot account for two opposite patterns, it must be making biased predictions among individuals with low and/or high levels of the m.3243A>G mutation. To test this supposition, we split the longitudinal dataset into low-and high-level subsets and evaluated the 2% annual decline model for bias in each of the two subsets. We noted, however, that while the analysis presented in Figures 1 and 2 does demonstrate the existence of dichotomy in mutation dynamics in blood, it does not permit precise determination of the threshold separating the two subsets with different mutation dynamics. To address this, we use 4 different thresholds (10, 15, 20, 30%) which essentially cover the entire span of the data to generate 8 subsets of the longitudinal dataset, 4 below the each of 4 thresholds (‘low mutant level subsets’) and 4 above the thresholds (‘high mutant level subsets’). The subset sizes are as follows: <10%, 18; <15%, 31; <20%, 39; < 30%, 61; >10%, 78; >15%, 65; >20%, 57; > 30%, 35; all, 96).). To test our hypothesis that the standard 2% annual decline (Rajasimha et al., 2008) model was unable to correctly handle some parts of the dichotomous data, we used the 2% model to predict mutant levels at the last measurement given mutant levels at the first measurement and the age difference between measurements. More specifically, we used the equation:

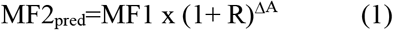

where R is the rate of change on mutant fraction per year, MF1 is the actual mutation level (MF stands for Mutant Fraction) at the first measurement, MF2_pred_ is the predicted level at the last measurement, and ΔA is the difference in Age between the two measurements. We then calculated the error ratio MF2_pred_/MF2 (where MF2 is the actual level at the last measurement) for each individual prediction and the geometric average of error ratios for each given subset (see also Figure 3A caption). Error ratios vary around 1, with 1 meaning ‘no error’. To make the measure of the error more intuitive, we then subtracted 1 from error ratios to obtain the ‘average relative error’ of the prediction, which is a fair measure of the bias of the model (positive or negative) in the given subset. The result of this analysis is shown in Figure 3A. As shown in forward predictions, the standard (2% annual decline) model underestimates mutation frequency (negative average relative prediction error) in the low mutant domain (blue) and overestimates (positive average relative prediction error) in the high fraction domain (red). This implies that the m.3243A>G decline rate of 2% per year is too slow for the high-frequency individuals, in whom mutations apparently decline on average faster than 2%, and too fast for the low fraction domain, where m.3243A>G level is more stable or increases with age within individuals on average.

**Fig 3:**
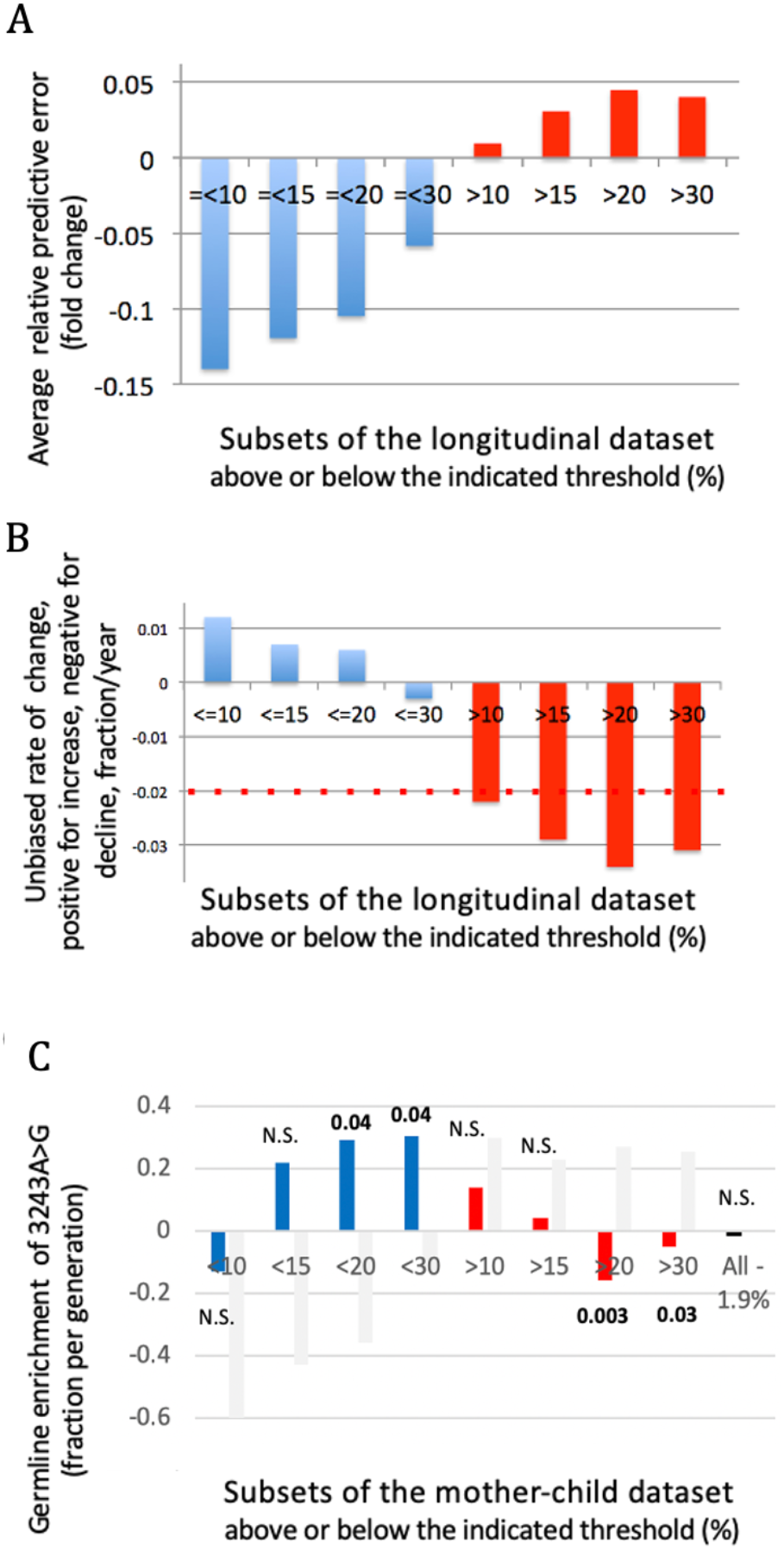
**(A) The standard 2% annual decline model, is negatively biased at low mutation levels and positively biased at high mutation levels.** Four high-mutation fraction (MF) and four low-MF subsets of the longitudinal dataset were created by splitting the dataset into two subsets at four arbitrary MF thresholds (10, 15, 20, and 30%; at first measurement). The 2% annual decline model was used to predict the level of m.3243A>G in each individual at last measurement based on the first measurement and age difference. The average relative errors within each subset were calculated and plotted as blue bars (low MF subsets) or red bars (high MF subsets). The colors were chosen to emphasise that high and low mutational level subsets preferentially consist of ascending and descending segments of Figs 1A and 2A. **(B) Subset-specific ‘unbiased’ mutation decline rates that neutralize biases depend on the mutational level of the subset**. ‘Unbiased’ rates were determined for each of 8 subsets described in (A), presented as the rates of annual decline R (equation (1)) such that error of predictions of the resulting ‘unbiased’ model averaged within the subset was zero. Blue bars – low-MF subsets, red bars – high-MF subsets. Red dotted line represents the conventional −0.02 (2% annual decline) rate. See Materials and Methods and Figs 4 and S1. **(C) ‘Unbiased’ model reveals positive germline selection**. To estimate ‘unbiased’ enrichment of m.3243A>G per generation due to germline selection, mother–child dataset was split into 8 overlapping subsets in the same way as longitudinal dataset in panel A. The unbiased rates derived from the 8 subsets of the longitudinal dataset shown in panel B were used to predict child’s mutational level at mother’s age and the ratio of adjusted child’s to mother’s mutation level was considered estimate of germline selection. Bars represent enrichments (i.e., geometric medians of child/mother ratios within each of the 8 subsets minus 1). Blue bars represent low-MF subsets, red bars – high-MF subsets. Black bar (‘All’) represents median enrichment (i.e., lack thereof) estimated by the standard 2% annual decline model within the entire mother-child dataset. Grey bars represent germline selection predicted by 2% decline model in each of the 8 subsets. Numbers above the bars are p-values (two tail sign test). See Materials and Methods for details.

### Biphasic models alleviate biases of the 2% annual decline model

To alleviate the biases of the standard 2% annual decline model, we propose to use a biphasic model which uses two annual decline rates, reflecting the dichotomous pattern of mutation decline with age. To build a biphasic model, the longitudinal dataset, which is used as ‘training’ data set, is divided into two subsets – below and above a chosen ‘separation threshold’ of the mutant fraction. For each of the two resulting subsets, an unbiased mutation decline rate is determined. This is the rate which, when used in equation (1), results in zero average error calculated over the subset (see materials and methods for details). The rationale is that if this model is unbiased in the training set, then it will most likely be unbiased in predicting future mutation levels on a blind dataset (like the mother/child dataset, where we need to predict mutation levels in children at their mother’s age).

This approach requires the specification of one free parameter: a threshold level of mutation fraction to separate the dataset into high and low mutation level subsets. The two ‘unbiased’ rates of annual decline of mutation level for each of the two subsets are then determined unambiguously by the condition that the model is unbiased. As far as the separating threshold is concerned, we used the same thresholds and generated the same eight subsets as those used for testing the conventional 2% annual decline model described in the previous section (Figure 3A and 3B). For each of the eight subsets, we determined the ‘unbiased’ rate of decline. Reassuringly, the ‘unbiased’ model for each subset was also close to the best fit model, meaning that the sum of squared errors was close to a minimum (see Supplementary Figure S1 for a full set of graphs).

Additionally, we showed that the biphasic model is significantly more unbiased than the standard 2% model. To prove this, we performed a simulation where the longitudinal dataset was randomly partitioned into equal sized training and testing sets multiple times (1000 iterations) and biphasic models were built for each drawing of the training set, and then tested on the test set. Then the performance of the biphasic models on the test sets was compared to performance of the standard 2% model. As expected, biphasic models performed unbiasedly on average, while the 2% model was biased, and the two distributions were significantly different (p<0.00001). The biphasic model was more unbiased than the 2% model in 70% of iterations. This statistic must be considered in light of the fact that this simulation was highly conservative, i.e., it was overwhelmed by random variation resulting from the small sample size: both training and test sets were much smaller than in real experiment (where the training set is the entire dataset from Grady et al. (2018), not half of it, and test set is the entire mother/child dataset, which is about 3-fold larger than the Grady et al. dataset).

As expected, the unbiased rates (Figure 3B) look like a mirror image of the average relative error bars (Figure 3A). Note that the line of reflection in Figure 3B is not the X-axis, but the red dotted line at −0.02, which is because the 2% annual decline model is the reference in this case. Interestingly, unbiased rates tend to be positive (i.e., mutant fraction increases in the blood with age) in individuals with a low mutant fraction, and negative in individuals with a high mutant fraction. This distinction remains relatively stable as the threshold separating the high and the low mutant fraction subsets is varied from 10 to 30%. The fair consistency of the neutral decline rates in subsets that lie predominately within the high (>15%, ‘>20%’,’>30%’) or within the low mutation level domain (‘<10%’, ‘<15%’, ‘<20%’) imply that the unbiased decline rates are innate characteristics of each domain. We therefore conclude that the high and the low mutation level domains should be analysed differently/separately. This is particularly true as the unbiased decline rates in the two domains have opposite signs, which are likely to compensate each other and obviate the details of the dynamics of m.3243A>G if they are (incorrectly) treated jointly.

### Estimating germline selection using the bi-phasic model

Germline selection of mtDNA pertains to the bias in the transmission of a mtDNA variant to the next generation, so ideally germline selection would simply be the ratio of mutation levels in the child vs mother. In reality, the child mother ratio is affected by random process of segregation of the two genotypes due to the intergenerational mtDNA bottleneck. As a result, selection can reliably estimated only by averaging the child/mother ratios among a substantial number of mother/child pairs. Geometric averaging which we use here is a natural way of averaging ratios. Furthermore, the child/mother ratio is inevitably based on m.3243A>G levels measured in a somatic tissue, most often, whole blood. Because m.3243A>G levels in blood systematically decrease with age, in the case of blood samples the child/mother ratio needs to be corrected for the age difference to yield an unbiased estimate of germline selection. We therefore expected that the bias of the 2% model that we described above might have affected previous estimates of germline selection.

To test this expectation and to obtain corrected estimates, we split the mother-child dataset 4-ways into eight subsets, mirroring our methodology for the longitudinal dataset. We then used, for each of the subsets, the specific unbiased decline rates devised from the longitudinal data (Figure 3B), to factor out the mother-child age differences within the corresponding subsets of the mother-child dataset. Children’s mutational levels were projected to mothers’ age using the ‘unbiased’ rate of m.3243A>G decline and divided by the mothers’ mutation levels. The details of the calculations for these estimates are presented in Materials and Methods. The estimated germline enrichment is positive at intermediate low mutant frequencies (<15%, <20%, <30% but not in the lowest subset <10% (Figure 3C). Notably, at higher mutant fractions positive selection decreases and then becomes negative. This suggests that m.3243A>G selection in the germline follows an ‘arched’ curve.

We then compared our results to the standard model: we calculated the expected germline selection for each of the subsets, and for the entire mother-child dataset, under assumption of the uniform decline rate of 2% per year. For the entire dataset, this produces a germline selection estimate which is close to zero (Figure 3C; black bar “All”). The subset-based analysis (Figure 3C; grey bars), unlike our bi-phasic approach which tends to predict weak positive selection estimates in most subsets, clearly predicts strong negative selection in the low mutation level domain and strong positive selection in the high mutation level domain. The explanation for this paradoxical behaviour of the standard approach is as follows: when a flat −2% rate is being used for age adjustment of the low mutation level subsets the mother-child pairs are being adjusted with a very incorrect mutation decline rate (2%/year, i.e., decrease instead of the unbiased 2%/year increase as in our calculations). This excessively negative rate results in excessive negative (instead of the needed positive) adjustment of the child’s mutation level projection to the mother’s age, which results in a very negative germline selection estimate. In contrast, in the high mutant fraction subsets, the use of decline rate of 2%/year is not sufficient to compensate for the even more rapid, and real, decline in these subsets. By our estimates, decline rates in high mutant subsets are about 3%/year, (average of the red bars in Figure 3C). Thus, insufficient negative adjustment leaves child’s levels too high, which results in an overestimate of positive germline selection.

Although our estimates of selection within the high m.3243A>G level subsets of the mother-child data are negative (the median-based estimate), the distributions are in fact skewed. While a majority of the data appear to show low levels of negative germline selection, there is also a long right-hand tail of what appears as positive selection (Supplementary Figure 2). The presence of these asymmetrically positioned outliers suggests that we cannot rule out the possibility of some positive selection taking place in these high-level subsets, potentially taking place in a special subset of individuals, which can be dependent, for example, on their nuclear genetic background.

## Discussion

### Dynamics of m.3243A>G in blood are dichotomous

Negative selection of the m.3243A>G pathogenic variant in human blood is well established; previous approaches to model this decline with age have pointed towards an exponential or sigmoidal process but the dynamics of m.3243A>G are more complex and these models do not fully-explain the data (Grady et al., 2018; Rajasimha et al., 2008; Veitia, 2018). Using a large quantity of recently compiled longitudinal data (Grady et al., 2018), we report that the dynamics of m.3243A>G in blood are dichotomous; levels predominantly decline in individuals with high mutation levels and are predominantly stable or even slightly increasing at low mutation levels. Therefore, the standard 2% annual decline correction, which is adequate on average, creates bias both at high and low mutation level and its predictions depend on the proportion of individuals with high and low mutation levels in the dataset. Interestingly, we detected a similar dichotomy in a large dataset of blood m.3243A>G levels in mother-child pairs, providing further support for a model in which the dynamics of m.3243A>G decline are dependent upon mutation level. Unbiased age correction is needed to circumvent the drawbacks of the 2% annual decline model; we used our observations to develop a new, empirical model of the decline of m.3243A>G in blood which better accounts for this dichotomy. We then used this unbiased age correction to explore the transmission of m.3243A>G from mother to child, detecting patterns consistent with positive germline selection of this variant, which may be a contributing factor in the comparatively high frequency of m.3243A>G compared to other pathogenic mtDNA point variations (Gorman et al., 2015).

Dichotomous dynamics cannot be described by a single decline rate. Therefore, any model that does not account for this dichotomy will be biased. To reveal this, we tested a 2% annual decline model as a predictor of subsequent measurements of mutation level based on preceding measurements and the age difference in different subsets of the longitudinal dataset. Average error is positive in the subsets of individuals with high mutation levels and negative in low mutation level individuals, confirming that the 2% annual decline model is biased. We note however, consistent with previous reports (Otten et al., 2018; Wilson et al., 2016), that it appears unbiased overall; the average error of its predictions across the entire longitudinal dataset is very small and not significantly different from zero. This can be explained by the neutralizing effect of the two partial biases, which occurs when averaging across both the high and low mutation subsets of the data. Interestingly, this means that overall bias of the 2% decline model is not universal – it depends on the relative composition of the dataset, i.e., how many individuals are in the high and in the low mutation groups.

### Biphasic model – an unbiased approach to dichotomous dynamics of m.3243A>G

Adjustment of the blood levels of m.3243A>G in mother and child for the age difference using the 2% annual decline model has been shown to have a dramatic effect on the estimates of germline selection of the m.3243A>G mutation as compared to unadjusted estimates (Rajasimha et al., 2008). We reasoned that this bias would significantly affect current estimates of selection as well. We therefore sought to modify the 2% model to eliminate this bias while keeping the elegant framework of the 2% model maximally unchanged. We chose to use a bi-phasic model to reflect the dichotomy of the m.3243A>G dynamics. In the interest of keeping the number of parameters in the model to a minimum, we made the simplest assumption that there are two subsets, i.e., the high and the low mutation level subsets, each with their specific exponential dynamics. It should, however, be noted that individual variability in decline rate still exists within these domains. We postulate that this could arise from differences in the threshold for biochemical expression of the variant allele between individuals, thus affecting negative selection against cells with high levels of the variant allele (see further discussion in the positive selection section). This theory is compatible with the high level of variability in disease burden and phenotypic spectrum that is seen in individuals carrying m.3243A>G (Chin et al., 2014; de Laat et al., 2012; Grady et al., 2018; Kaufmann et al., 2011; Mancuso et al., 2014; Nesbitt et al., 2013; Pickett et al., 2018). Additionally, our preliminary studies imply that that the variability of the decline rates between individuals may in part result from complex dynamics of the cell composition of whole blood, where different cell types may carry different mutational loads, according with a recent report (Walker et al., 2020).

In the initial stages of this study, we attempted to use complex models with extra parameters to capture the variability of the data and were convinced that given a relatively small size and high variance of the dataset, such attempts resulted in overfitting. We have therefore chosen to limit analysis to the most basic model possible for a dichotomous data – the biphasic model.

The development of this model revealed important features of the dynamics of the mutation in blood. Contrary to the conventional view that blood m.3243A>G levels universally decrease with age, in the blood of individuals with low mutation levels, m.3243A>G levels tend to stabilise and even increase in some cases. Conversely, unbiased rates in the high mutation level individuals are more aggressively negative than conventionally accepted decline rate of 2%. Thus, the perceived overall 2% decline rate of the standard model probably stems from de facto ‘averaging’ of the rates at high and low mutational levels, which has been possible to detect due to the recent availability of larger, longitudinal datasets.

### Analogy between the longitudinal and the mother/child datasets

The similarity that we see between the longitudinal and mother-child datasets is expected and thus is reassuring. Indeed, m.3243A>G levels in child and mother can be viewed as two sequential samplings of mtDNA mutation levels from the same germline. By ‘sampling’, we mean that the continuous germline gave rise to somatic tissue (blood cells in our case) in the mother and then in the child, each of which were used to infer mutation level of the germline. These two samplings, however, systematically differ in that in the child the mutation spends fewer years in the blood cells than in the mother. Similarly, in the longitudinal dataset, every pair of sequential measurements represent two samplings of the germline.

### Unbiased bi-phasic approach reveals positive selection in the germline

Germline selection of m.3243A>G can be estimated by comparing mutation levels in children and mothers. However, because of the systematic changes in the blood levels of m.3243A>G with age (Rajasimha et al., 2008), the relative levels in blood in mothers and children must be adjusted for the age difference. Previously, this has been achieved by applying the annual decline model of ∼2% per year (Wilson et al., 2016). We revisit the transmission dynamics of m.3243A>G using the bi-phasic model developed in this study. Accordingly, we divided mother-child pairs into a series of high and low mutation level subsets and used the subset-specific unbiased rates as the best empirical means to correct for the mother-child age difference needed for estimation on intergenerational germline selection. Results of this analysis are shown in Figure 3C. We do note a variability in the estimates; obtaining more precise estimates of the magnitude of this apparent germline selection will require larger and more balanced datasets.

The distribution of positive selection estimates shown in Figure 3C implies that positive selection is strongest and most consistent in subsets which include cases of moderately low mutant fraction levels (<20, <30). Currently it is difficult to precisely determine the extent of selection at lowest mutant fractions (i.e. when MF approaches zero). We note however, that our analysis provides no evidence that positive selection persists at lowest mutant fractions, rather it most likely decreases, converges to zero or even becomes negative. This conclusion is corroborated by the apparent shift of the main peak of the distribution to the zero position (compare distributions ‘<20’, ‘<15’ and ‘<10’ in Supplementary Figure 2). More data are needed to substantiate this preliminary conclusion.

We also saw asymmetric outliers within the higher-level subsets; therefore, we cannot discount the possibility that some positive selection is also occurring at higher levels. As all of these outliers are so dramatic, it is tempting to speculate that excessive individual variability of decline rates may have genetic cause. Indeed, familial clustering was demonstrated in a previous study which showed a high heritability estimate for m.3243A>G level (h2=0.72, standard error=0.26, P=0.010) (Pickett et al., 2019). Moreover, another recent study has identified several nuclear genetic loci associated with non-pathogenic heteroplasmy (Nandakumar et al., 2021). Further study of the effects of nuclear genetics on the transmission of pathogenic mtDNA variants is warranted.

What can be the mechanism behind positive selection? It is tempting to speculate that, for example, m.3243A>G may offer a positive advantage – potentially because less efficient translation of mtDNA-encoded proteins may cause local compensatory mtDNA replication or stimulate cell proliferation (Smith et al., 2020), both of which will result in positive germline selection. At higher mutation levels, however, translation deficiency is expected to result in progressive respiration defect that is likely to eventually trigger oocyte attrition, or embryo demise, as has been demonstrated for other detrimental mtDNA mutations, e.g., (Fan et al., 2008; Freyer et al., 2012). This effect would explain the trend towards negative selection at higher mutant fractions seen in Figure 3C.

### Bi-Phasic vs. 2% annual decline approach: the sources of the discrepancy

To put these findings in the context of previous studies, we performed additional analysis to explore the sources of differences between our estimates and those reported previously, which found no evidence of positive selection (Otten et al., 2018; Wilson et al., 2016). First, we directly compared our approach to the 2% annual decline model by applying this correction to exactly the same mother-child dataset that we used for the bi-phasic estimates, to exclude the possibility that differences are caused by differences in the dataset. Reassuringly, in agreement with previous studies, the predicted overall germline selection when calculated over the entire mother-child dataset using the standard 2% decline rate is very low and not significantly different from zero. Most notably, unlike our biphasic approach, which tends to predict weak positive selection estimates in most subsets, the 2% annual decline approach clearly predicts strong negative selection in the low mutation level domain, and strong positive selection in the high mutation level domain, an effect that we believe is due to insufficient adjustment in the high m.3243A>G level domain and excessive adjustment in the low m.3243A>G level domain.

The above observations reveal an important drawback of the standard model. Estimates based on the 2% annual decline model are highly sensitive to the composition of the mother-child dataset. Indeed, from Figure 3C (grey bars) one can deduce that the higher the proportion of low mutation level mother-child pairs included in the dataset, the lower the expected overall estimate of the germline selection using 2% adjustment. In fact, the near zero estimate of selection intensity in the current mother-child dataset is merely the result of the particular proportion of low and high mutation level mother/child pairs in this dataset. The biphasic model is poised to deliver much higher stability of the estimate with respect to the changing proportion of high and low mutation level patients.

### Comparison to other studies: positive and negative germline selection

Convincing positive germline selection of the m.3243A>G has not been reported previously but has been observed for other detrimental mtDNA variants. An apparently milder variant, m.8993T>G, may be under positive germline selection throughout the mutant fraction range (Otten et al., 2018). Interestingly, data from Freyer et al. (2012) (Freyer et al., 2012) show that a specific highly detrimental mutation, the m.3875delC in the mouse mt-Tm gene, encoding tRNA^Met^, at an intermediate mutant fraction (45-60%) systematically increases its presence in offspring compared to the mother, implying positive selection of a detrimental mutation in the mouse germline which is reversed at higher (>60%) levels (see their Figure 2A and Figure 3). Surprisingly, Freyer et al. did not discuss their own data as supporting positive germline selection. Of note, no data are available for mutational levels below 45% for this mutation, so it is possible that selection decreases at lower mutant fraction as we observe for m.3243A>G (Figure 3C). In conclusion, it appears that detrimental mutations, at least in some cases, are under positive selection in the germline. It would be interesting to determine how general such a phenomenon is, using a broader set of detrimental mtDNA mutations.

A few months after original version of this paper was first published in BioRxiv (Fleischmann et al., 2021), another group has reported positive germline selection of the m.3243A>G (Zhang et al., 2021).They reported positive selection, but their conclusion was based on an incorrect analysis. Specifically, they showed that positive selection (presented as average heteroplasmy shift, HS) was highest at low mother mutation frequency values, gradually decreased with mutation frequency and eventually became negative at high frequencies. Our preliminary analysis indicates that discrepancy between our results (arching selection profile) and those of Zhang et al. (monotonous decrease of selection with mutation level) can be accounted for by the ‘regression to the mean’ bias (Galton, 1889), which adds strong spurious negative correlation between the shift of mutant fraction from mother to child and mother’s mutant fraction.

### Potential applications of the biphasic approach

From a practical point of view, this study will hopefully lead to models that better account for the dynamics of pathogenic mtDNA variation by including the effect of mutation level. The simple biphasic model proposed here may serve as a practical alternative to the 2% annual decline model in cases where, like in studies of germline selection, unbiased prediction and versatility of the model (applicability to datasets that are unbalanced with respect to cases with low and/or high mutational levels) is essential. This can be realized by applying one of the two rates: ∼3% annual decline for cases with mutation levels above 20% and ∼1% annual increase for cases with levels below 20%. Of note, in this study we used the mutation level at first measurement, or mutation level of the child (which is analogous to the first measurement). We have chosen this convention because the dynamics of the mutation converges with time (phase lines become denser with time), so, in general, prediction of the succeeding mutation level is more stable than prediction of the preceding mutation level. As of now, this bi-phasic model is not a finished working instrument that can be used to estimate at-birth m.3243A>G levels within a clinical prognostic setting. Rather, this research reveals that dynamics of m.3243A>G in blood is more complex than previously considered and highlights the need for more detailed, perhaps mechanistic, models, especially in the lower range of mutation levels. With more data available for models to build upon and test, our approach may be further optimized or replaced by a more detailed and precise model. Our current model, however, is appropriate to make conclusions about the aggregate behaviour of mutations, such as positive selection, with the data currently at hand.

## Materials and Methods

### Data sources

Longitudinal measurements of m.3243A>G levels in blood were obtained from Grady et al. (Grady et al., 2018). This dataset is comprised of 96 individuals who were recruited into the Mitochondrial Disease Patient Cohort UK and had two or more measurements of m.3243A>G blood level taken between 2000 and 2017. The median time between first and last measurement was 2.5 years (IQR = 4.35, range = 0.01 – 15.20), the median age at first measurement was 36.25 years (IQR = 20.58, range = 15.60 – 72.50) and at last measurement was 40.25 years (IQR = 21.00, range = 19.80 – 78.80).

### Mother-child data

Measurements of m.3243A>G levels in mother-child pairs were obtained from Pickett et al. (Pickett et al., 2019). This dataset contains 183 mother-child pairs (from 113 different mothers), comprised of 67 pairs from the Mitochondrial Disease Patient Cohort UK (Nesbitt et al., 2013) and 116 pairs obtained from a literature search and previously published by Wilson and colleagues (Wilson et al., 2016). To minimize ascertainment bias, pairs in which the child was the proband had previously been removed, as had one pair from the literature where the m.3243A>G variant was thought to have arisen de novo in the child (Ko et al., 2001). We identified an additional 30 pairs where the mother was the proband; to reduce bias these were also excluded, leaving a total of 153 pairs (from 97 mothers) that were taken forward into the analysis. The mean age at m.3243A>G level assessment was 53.9 years (SD = 12.4, range = 23.0, 85.0) for the mothers and 27.8 years (SD = 12.2, range = 0.4, 58.0) for the children.

### Construction of Binary Logistic Regressions

The binary logistic regressions were performed on both the longitudinal and mother-child data sets. For each data set, the final mutation level was subtracted from the initial mutation level. Then, each pair of datapoints were assigned a binary value for their mutation level directional change. A “1” was assigned for increasing mutation levels and a “0” was assigned for decreasing mutation levels. Data point pairs that had no change were excluded from the analysis. A simple binary logistic regression was performed on each set of data using the initial mutation level values of each data point pair as the independent variable and the corresponding binary indicator as the dependent variable. Each regression was performed with a likelihood ratio test, goodness of fit test, and with 95% confidence intervals. Significance was recorded as p-values. For each binary regression for each data set, the predictive curve was plotted with the respective data points. The longitudinal data set was graphed individually (Figure 1B), and the longitudinal and mother-child data set was graphed together for comparative analyses (Figure 2B). Statistical analyses and graphing were performed in GraphPad Prism version 9.

### Determination of the unbiased mutation decline rates

The approach we used to calculate the unbiased mutation decline rates in the various subsets of the longitudinal dataset (Grady et al., 2018) is illustrated in Figure 4. Of note, we limited analysis to the first and the last measurement for each person, so that there were only two measurements for each person in the dataset, mf1 and mf2 (‘mf’, for mutant fraction). For every individual (of 96 individuals in the dataset) and thus every pair of data points (mf1, mf2), the mf2 was predicted based on mf1 using an exponential model with variable parameter R, i.e., the fractional decline of mutational load per year (R is positive for an increase, and negative for decline) where ΔA is the age difference between the times of mutation level measurements. This is function (1) as described in the results section. R was varied between 0.05 decrease to 0.05 increase per year (i.e., from −0.05 to 0.05) as shown in Figure 4 in steps of 0.0000001 (1,000,000 steps overall). Thus, we obtained 1,000,000 values of mf2_pred_ (R) for each R, for each individual (i.e., 100×96 total). For each mf2_pred_ (R), an error ratio (mf2_pred_ (R)/mf2) and absolute error (sqrt[(mf2pred/mf2)]^2^) was calculated and then both error and absolute error were averaged among individuals within each of 8 subsets. We obtained 1,000,000×2×8 of the averaged data points and plotted them for each value of R in 8 graphs shown in Supplementary Figure S1 (one graph per subset). The minimum of the absolute error (corresponding to the best fit rate) and the zero of the average non-absolute error (“unbiased rate”) were determined graphically by identifying the minima of the curve and the intercept of the x axis, respectively. Unbiased rates were then used for plotting the graphs in Figure 3B. The best fit rates were used to make sure that the unbiased model was close to the best fit model.

**Fig. 4:**
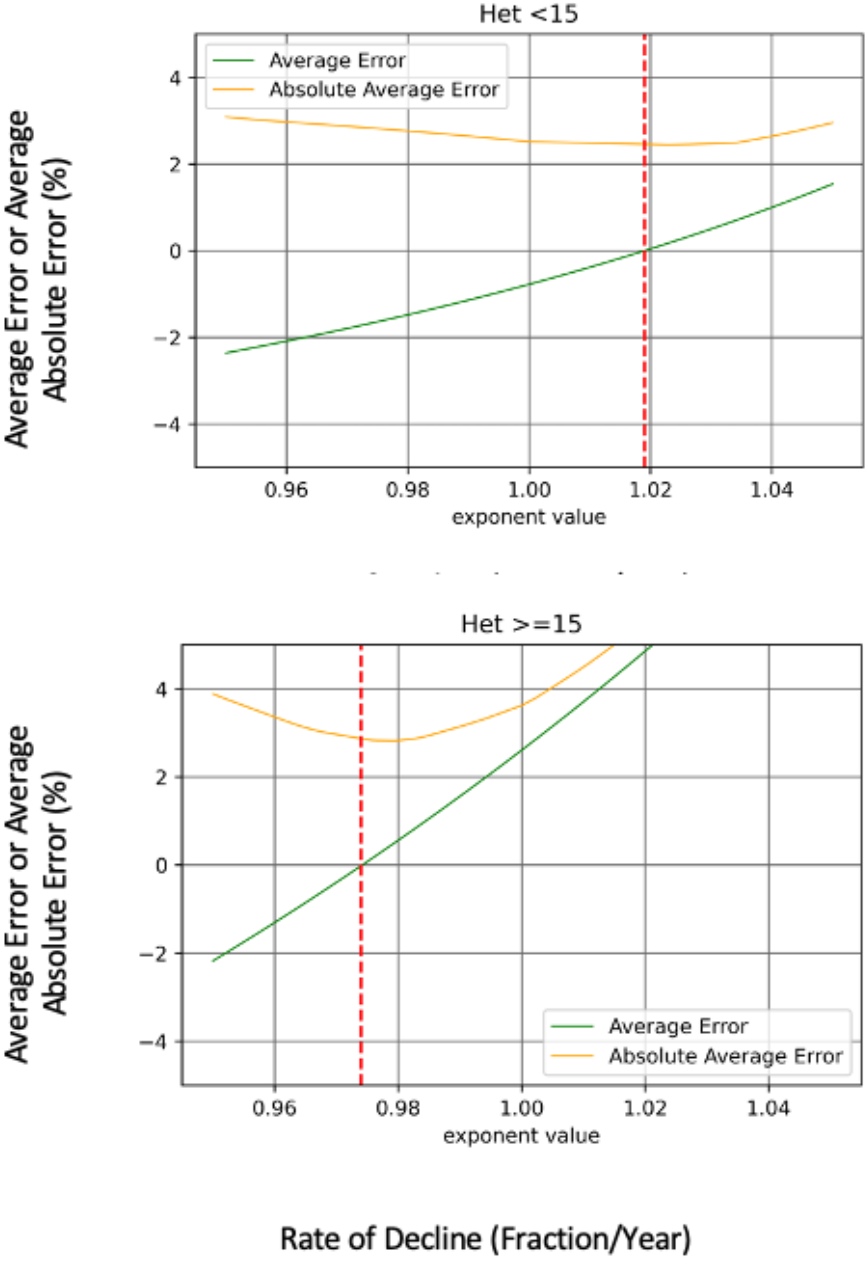
Example of the calculation of an unbiased rate of decline for the two subsets: <15% (top) >=15% (bottom). See Materials and Methods for the procedure and Supplementary Fig. S1 for a complete set of curves. Red vertical lines indicate the unbiased rate.

### Calculation of the enrichments/selection of m.3243A>G per generation in the germline

The enrichments/selection per generation for a specific subset of the data S is determined as geometric average of ratios of age-corrected child mutation levels to those of their mothers. In practice, we calculated the average of logarithms of the ratios and then converted the average back from logarithm to real ratio/geometric mean. That is, we first determine the germline selection for each child-mother pair within each subset using Equation 2 –

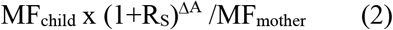

where R_s_ is the estimated unbiased rate for the subset S to which the mother-child pair has been assigned (as determined by the value of MF_child_) and both MF_child_ and MF_mother_ are expressed as proportions between 0 and 1. ΔA is the age difference between mother and child.

For example, for the 20% threshold, a child with a m.3243A>G mutation level of 40% would be assigned to the ‘>20%’ subset with an R_s_ of −0.029 (see Figure 3A). If their mother was 30 years older and had a m.3243A>G level of 10%, the germline selection estimate in that individual mother/child pair would be: 0.4 × 0.971^30^ / 0.1 = 1.65

We then calculated median log ratios across all mother-child pairs in the given data subset and used two-tailed sign test to determine the p-value associated with this median of logarithms being different from zero (rejection of the null hypothesis of no selection). Finally, the median log ratios for the various data subsets were converted into ‘median mutation level ratios’ by calculating exponent of the median of log ratios and 1 was subtracted from it to produce “median enrichment” (i.e., fractional increase, positive for enrichment, negative for depletion, analogous to the rate of decline used describe the dynamics of m.3243A>G in blood). Kernel density estimation was performed using on-line portal at http://www.wessa.net/rwasp_density.wasp with default parameters.

## Conflict of Interest Statement

The authors declare no conflict of interest.

### Supplemental Figures

after the Reference list.

---

**Supplementary Figure S1.**
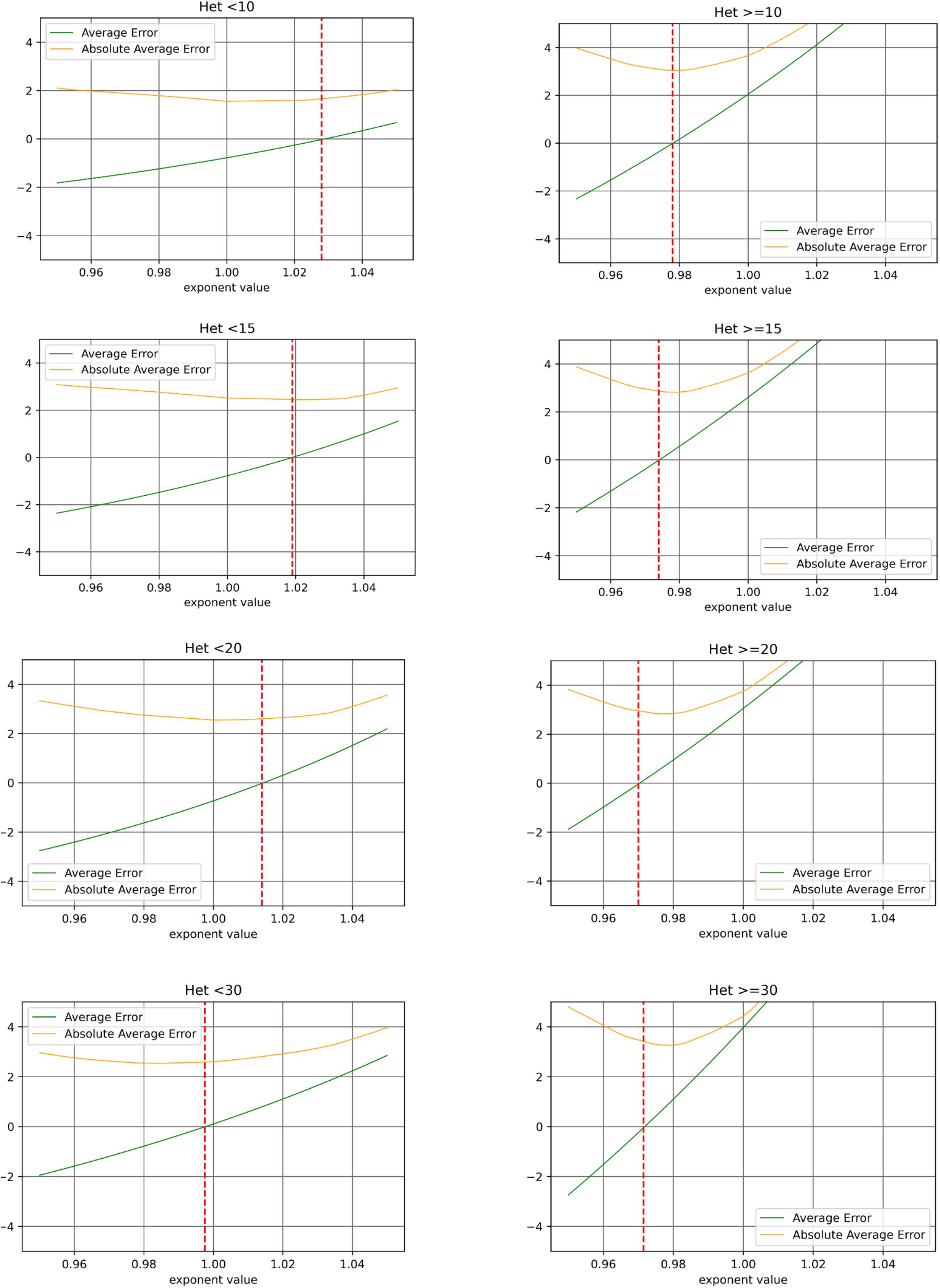
Full set of graphs used to determine the unbiased decline rates.

**Supplementary Figure S2.**
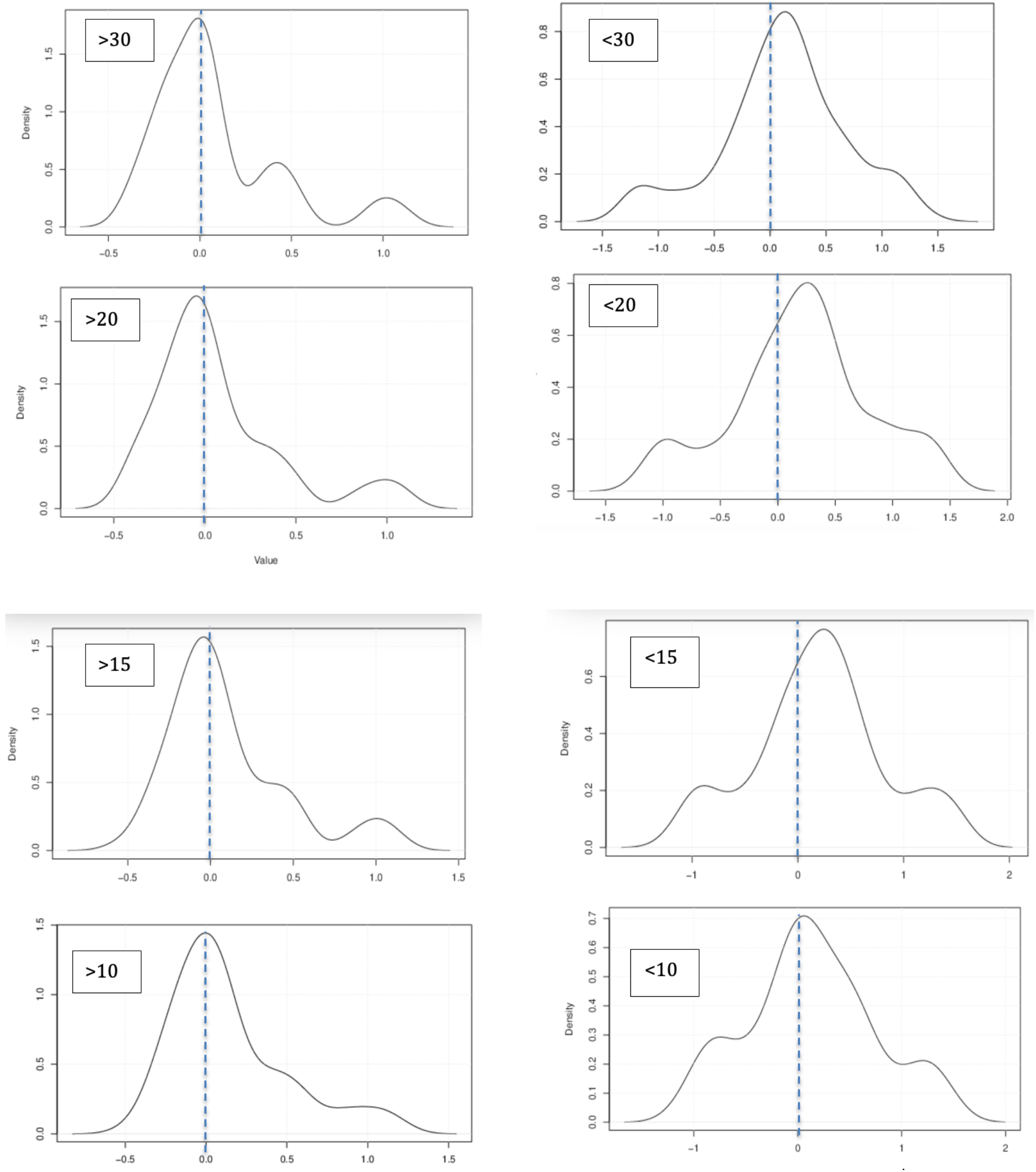
Images of the kernel histograms of the age-corrected shifts between mother and child.

## Notes

This study was supported by a grant from the U.S. National Institutes of Health (R01-HD091439 to J.L.T., D.C.W. and K.K.), KP was supported by the Ministry of Science and Higher Education of the Russian Federation (agreement no. 075-02-2022-872), Russian Science Foundation grant 21-75-20143 and Russian Federal Academic Leadership Program Priority 2030 at the Immanuel Kant Baltic Federal University. This research was funded in part, by the Wellcome Trust [204709/Z/16/Z to S.J.P. and 203105/Z/16/Z to the Wellcome Centre for Mitochondrial Research]. For the purpose of Open Access, the author has applied a CC BY public copyright licence to any Author Accepted Manuscript version arising from this submission. We would like to acknowledge those involved in the collection of data for the Grady et al. (Grady et al., 2018) and Pickett et al. (Pickett et al., 2019) manuscripts.

### Competing Interest Statement

The authors have declared no competing interest.

